# The Perceptual Shaping of Anticipatory Actions

**DOI:** 10.1101/184333

**Authors:** Giovanni Maffei, Ivan Herreros, Marti Sanchez-Fibla, Karl J. Friston, Paul F.M.J. Verschure

**Affiliations:** Laboratory of Synthetic, Perceptive, emotive and Cognitive Systems (SPECS), Universitat Pompeu Fabra (UPF), Barcelona, Spain; Imaging Neuroscience and Theoretical Neurobiology, Wellcome Trust Centre for Neuroimaging, University College of London, London, UK; Institucio Catalana de Recerca i Estudis Avancats (ICREA), Barcelona, Spain; Institute for Bioengineering of Catalonia, Barcelona, Spain

**Author notes:** These authors equally contributed to this work (Giovanni Maffei, Ivan Herreros).

## Abstract

Humans display anticipatory motor responses to minimize the adverse effects of predictable perturbations. A widely accepted explanation for this behavior relies on the notion of an inverse model that, learning from motor errors, anticipates corrective responses. Here, we propose and validate the alternative hypothesis that anticipatory control can be realized through a cascade of purely sensory predictions that drive the motor system, reflecting the causal sequence of the perceptual events preceding the error. We compare both hypotheses in a simulated anticipatory postural adjustment task. We observe that adaptation in the sensory domain, but not in the motor one, supports the robust and generalizable anticipatory control characteristic of biological systems. Our proposal unites the neurobiology of the cerebellum with the theory of active inference and provides a concrete implementation of its core tenets with great relevance both to our understanding of biological control systems and, possibly, to their emulation in complex artefacts.

## 1 INTRODUCTION

Trained snowboarders smoothly control the board while adjusting their posture to anticipate bumps on the slope. Otherwise, at the speed of their descent, reacting only after the effects of the irregularities on the terrain are felt would cause them to lose balance and fall. Anticipatory motor actions, thought to depend on the cerebellum (1,2), are quintessential for skilled performance in sport, but they are also part of our everyday behavior: from walking (3–5), to grasping (6–8) and to riding a bicycle (9). The question then arises as to how these actions are controlled? Decades of research in motor control support the notion that internal models are key to skillful performance (10–12). Specifically, this research has highlighted two kinds of internal models: forward models, which map the efference copies of motor commands into their expected sensory consequences (13,14); and inverse models, which map desired sensory outcomes into their required motor commands (15,16).

However, here we argue that offering an alternative to these interpretations is a pressing issue for the field of motor control as neither forward nor inverse models (in their standard formulation) can explain the versatile anticipatory control observed in animals. In particular, standard forward models allow for rapid feedback control in the presence of the long transport latencies of the nervous system (13,17) or action planning (18) but, as they exclusively predict the consequences of motor commands, they cannot anticipate disturbances that are not contingent upon those motor commands (19). That is, one cannot call upon efference-driven forward models to support behaviors that precede external events. This obvious limitation has led researchers to conclude that preparatory actions should result from inverse models that output anticipatory motor signals (20–25). The benchmark computational model for that theory is feedback error learning (FEL), which offers both an adaptive motor control architecture (26) and a theory of cerebellar functions (10,16). In FEL, *predictive* actions are the result of anticipatory motor signals, learned by shifting forward in time the output of the feedback controller (20,25). However, we will show that inverse model schemes present some important limitations in the context of anticipatory control. For instance, while rapid corrections of erroneous anticipatory actions are commonly reported in biological systems, most notably in experiments that include catch trials (i.e. trials where a predictable disturbance is signaled but not delivered) (27), FEL has no mechanism to correct feed-forward motor responses once the course of events violates a prediction. In addition, FEL acquires motor commands that are tied to the dynamics of the plant that it controls and cannot easily be generalized to new configurations. However, experimental evidence suggests that in humans anticipatory responses are still effective even if one changes the posture and/or the effector after learning (28–30). Hence, given that standard forward and inverse models cannot fully account for anticipatory control, alternatives should be considered that both overcome the theoretical and practical limitations of these motor-centric accounts and resolve the forward-inverse model dichotomy.

Here, we advance the hypothesis that biological anticipatory control can be explained by the ability of the brain to advance predictions of future perceptual events (31,32) and use those predictions to drive the motor system in an anticipatory way (33). We formulate this hypothesis in computational terms by proposing the cerebellar-based Hierarchical Sensory Predictive Control (HSPC) architecture, in which internal models issue sensory predictions that facilitate anticipatory control, with motor signals (i.e. efference copies of motor commands) playing no role in adaptation itself. With that, HSPC challenges the inverse model interpretation of anticipatory control – and, indirectly, the “motor centric” forward-inverse model dichotomy. More precisely, we suggest that, in contrast to the FEL hypothesis, where predictive actions are the result of anticipatory motor signals, anticipatory actions can be controlled by predictive sensory signals, becoming reactions to events that are brought forward in time (34–36). Moreover, in HSPC the internal generation of sensory predictions can mirror the causal structure of the sequence of perceptual events (Fig. 1). HSPC builds on the theory that motor control can be understood as a process that deals with sensory predictions that are only mapped onto motor commands at the latest stage, e.g., by reflexes (37–39). This view has been studied mostly at the theoretical level, within active inference, using the formalism of generative hierarchical models and focusing on the aspect of reformulating control as Bayesian inference (40,41). Hence, to the best of our knowledge, here we propose the first computational treatment of this theory in the context of anticipatory actions. To this end we provide a systematic comparison between HSPC and FEL by synthetizing each hypothesis into an architecture applied to a postural control task, minimally modeled as the stabilization of an inverted pendulum through a torque at its base (i.e., ankles) (Fig. 1), demonstrating how learning in the sensory rather than in the motor domain can account for the robustness and generalization capabilities of biological control systems. In summary, the present study presents an approach to motor control that could provide an alternative interpretation of the physiology of anticipatory control and contribute to the theory of cerebellar learning.

**Fig. 1-.**
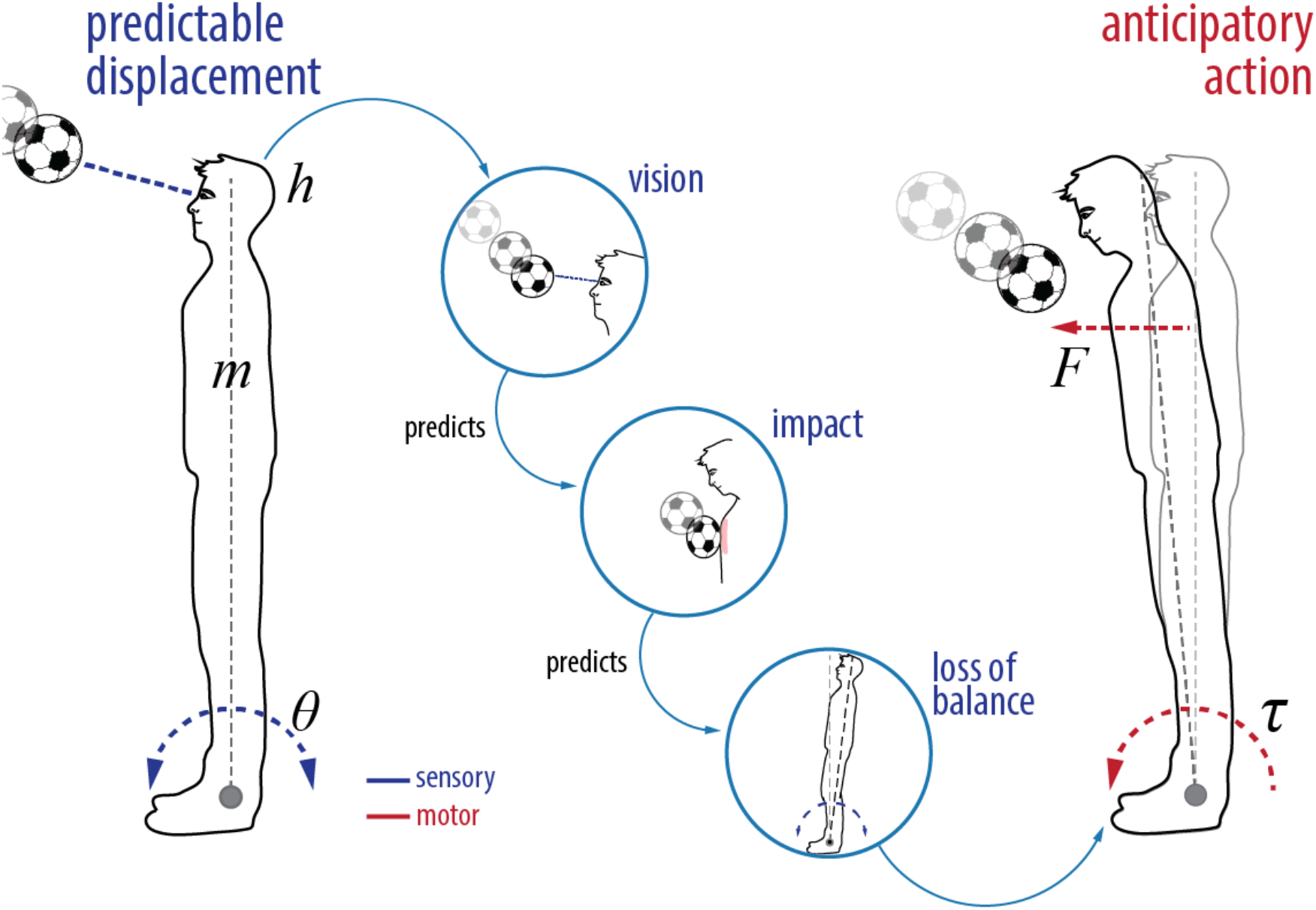
Conceptualization of the HSPC hypothesis. A predictable displacement caused by a soccer ball directed to the chest elicits an anticipatory response that minimizes the loss of balance before it is perceived. In HSPC the anticipatory response is the result of a hierarchy of descending sensory predictions from distal (visual detection) to proprioceptive (impact) to vestibular (loss of balance) modality, where each modality advances in time the expected consequences on the next modality until the predicted error in balance triggers a reflexive action in a feed-forward manner. The minimal model for this behavior is an inverted pendulum of mass (*m*) and height (*h*), whose error in angle (*theta*) is minimized by generating a torque (τ) at the ankles.

## 2 METHODS

In order to compare the behavior of a control strategy based on motor anticipation (FEL) with one based on sensory prediction (HSPC) we synthetize these hypotheses into two architectures that control an inverted pendulum (a common model for bipedal postural control–see (42) for review) (Fig. 2-B and 2-C) engaged in an anticipatory postural adjustment (APA) task. This task, in line with experimental psychology paradigms (2,43) (Fig. 2-A), requires the agent to learn an appropriate combination of anticipatory and compensatory responses to minimize the effect of a disturbance (i.e. loss of balance) signaled by a cue.

**Fig. 2-.**
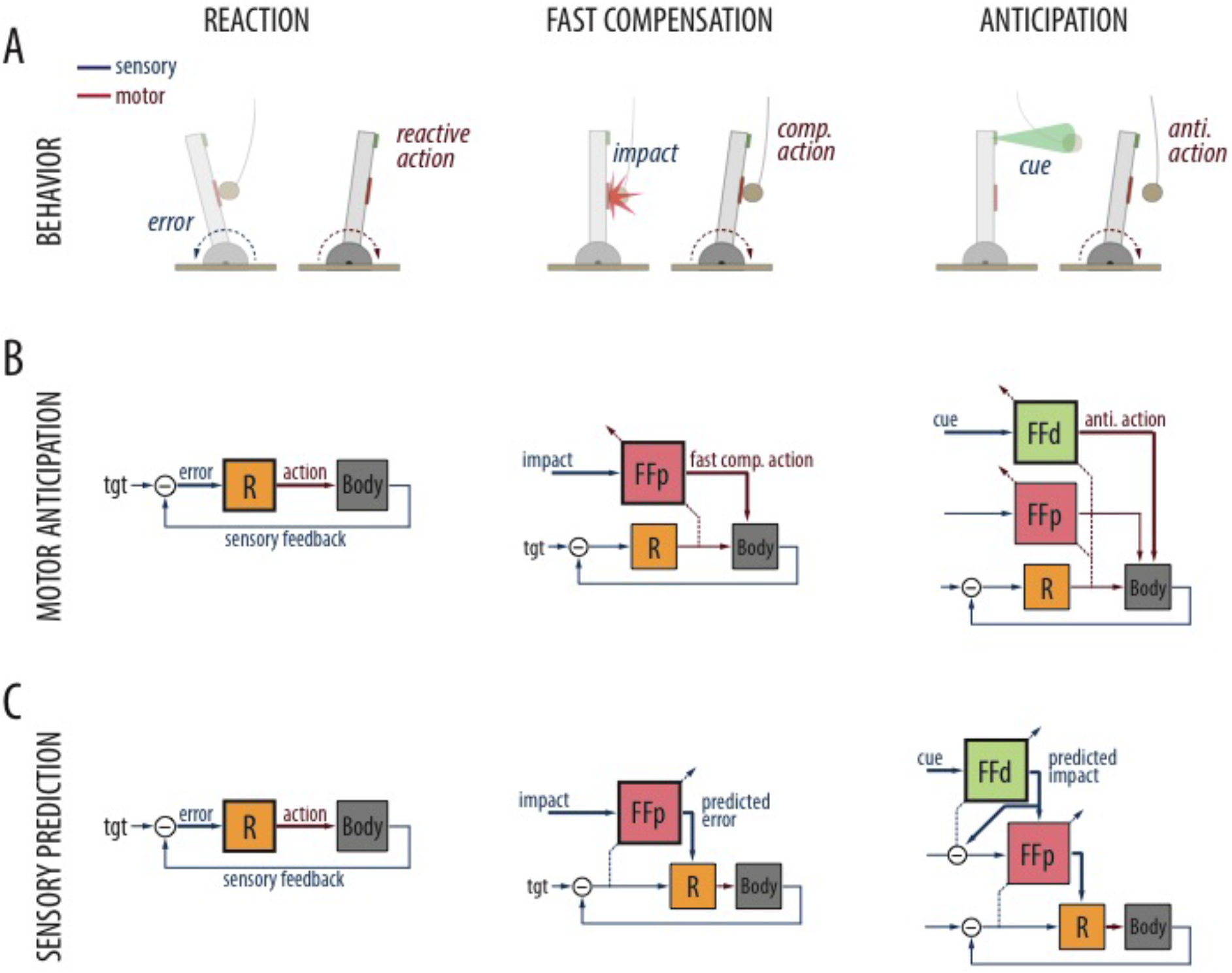
Motor anticipation (FEL) vs. Sensory Prediction (HSPC) strategies. **A.** Different responses are elicited by different sensory modalities. (From left to right): A corrective reaction is triggered by the perceived postural error. A fast compensatory corrective action is triggered by the perceived impact (proximal stimulus). An anticipatory action is triggered by the distance to the obstacle (distal cue). **B. Motor anticipation strategy (FEL):** A postural error is converted into a reflexive action by a feedback controller (R). A feedforward compensatory action associated to the impact signal is acquired by the proximal adaptive module (FFp) on the basis of the feedback response to error. A feedforward anticipatory action associated to the distal cue is acquired by the cerebellar distal module (FFd). **C. Sensory prediction strategy (HSPC)**. Reflexive action elicited as in FEL. Fast compensatory action: Triggered by the proximal cue and learned from the closed-loop error, a counter-factual error is issued by the proximal module (FFp) in response to the proximal cue driving the feedback controller. Anticipatory actions: Evoked by the cue, a prediction of the expected impact issued by the distal module (FFd) triggers the compensatory response in an anticipatory manner.

### 2.1 Model of the agent

The inverted pendulum actuated by a torque (*τ*) at its base is modeled as follows:

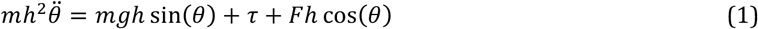

The pendulum has a mass (*m*) of 67 Kg and a height of its center of mass (*h*) equal to (0.85 m). *θ* measures the angular deviation from the vertical position. The disturbance is introduced as a force (*F*) parallel to the ground applied to the center of mass.

### 2.2 Control Architectures

The APA task involves three different sensory modalities: *distal exteroception* (perceiving a cue that precedes the collision), *proximal exteroception* (that could play the role of proprioception in humans, sensing the magnitude of the impact on the body) and *interoception* (sensing the postural effects of the impact, that is, the inclination). Each modality enables a different type of response: distal sensing allows for preparatory responses, proximal sensing for fast compensation and interoception for compensation through feedback control (6,34,44–47) (Fig. 2-A).

#### 2.2.1 Feedback controller

The agent is stabilized by a torque generated through Proportional-Derivative (PD) feedback controller as follows:

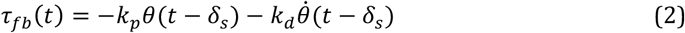

Note that in the error term we use the angle and angular velocity values delayed by *δ_s_* (= 100 ms.) to account for the latency of the error feedback.

#### 2.2.2 Adaptive feed-forward modules

In addition to a feedback controller, both architectures include the same adaptive feed-forward modules to process the proximal and distal cues. That adaptive feed-forward module (i.e., inversion of a forward or generative model) is implemented as an adaptive filter extended with an eligibility trace mechanism (48–50). Each feed-forward module receives a single (exteroceptive) input signal that is expanded into N (=20) different signals or bases. Each basis corresponds to the convolution of the (sensory) input with an alpha signal that can be formulated as two serially linked leaky integrators with identical time constants. For a particular basis, its output value is generated as follows:

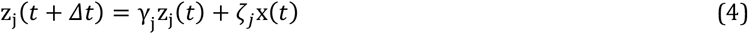

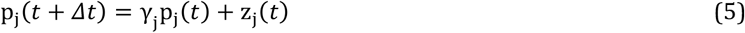

Where *Δt* (= 0.01 *s*) is the simulation time step and γ_j_ = e^-τ_j_Δt^ is the j-th basis decay factor, derived from a relaxation time constant τ_j_. ζ_j_ is a scaling factor that equalizes the power of all bases. At this point, an expansion of the original signal x(t) into a series of bases or transients with different temporal profiles obtains. The second processing step consists in mixing those bases according to a weight vector ***w*** (*t*) to generate an output signal (*ff*(*t*)):

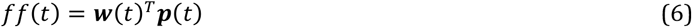

Where ***p***(*t*) = [*px*(*t*),…, *p_N_*(0]^T^ is the vector of the bases. The weight vector is adaptively set by means of a LMS or Widrow-Hoff update rule (51) extended with an eligibility trace:

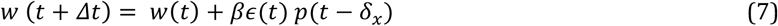

Here, *ϵ*(*t*) is an appropriated error signal that is used to update the weights. The eligibility trace is implicit in the use of a delayed copy of the bases activity *p*(*t* — *δ*_*x*_) for the update, with *x* indexing the type of stimulus processed: proximal (*p*) or distal (*d*). In short, to update the weights we associate the current error with an activity on the basis signals *δ_x_* seconds ago. With that we assume that activity at time *t — S_x_* is the activity that should have been used to trigger a reaction with sufficient anticipation to cancel the current error. In general we set both *δ_d_* and *δ_p_* greater than the error feedback delay (*δ_s_*), implying that the extent of the anticipation goes beyond the transport delay.

#### 2.2.3 Configuration of the FEL and HSPC architectures

Both control architectures include the feedback controller and two feed-forward modules (distal and proximal) wired according to the heuristic of either predicting motor commands from sensory signals (FEL architecture), or predicting sensory signals from sensory signals (HSPC architecture).

In FEL, feed-forward modules act upon the plant and are supervised by the feedback reaction to the error in posture (Fig. 2-B). In particular, the proximal module issues a feed-forward action in response to the impact learned by shifting the reactive action earlier in time, while the distal module similarly acquires a response that is triggered by the distal stimulus, and thus can precede the impact itself.

Let *ff_p_*(*t*) and *ff_d_*(*t*) be the outputs of the proximal and distal feed-forward modules; *x_p_*(*t*) and *x_d_*(*t*), their respective input signals; and *ϵ_p_*(*t*) and *ϵ_d_*(*t*) their respective teaching signals. The structure of the FEL architecture is determined by the following equations:

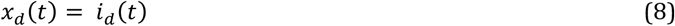

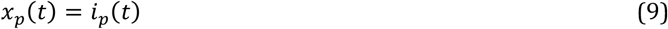

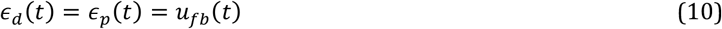

Where *i_d_*(*t*) and *i_p_*(*t*) represent the cue (*distal*) and impact (*proximal*) signals, respectively, and *u_fb_* is the output of the feedback controller. As a final step, the output of all modules are added up to generate the control signal (*u_fel_*(*t*)):

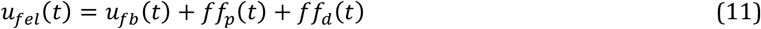

In HSPC, upstream modules drive and learn from the input of downstream modules (Fig. 2-C). That is, the proximal module learns *counterfactual* errors (33) contingent to the impact so that the feedback controller reacts to the expected error before the actual one occurs. While the distal module learns to predict the collision signal contingent to the cue and triggers the proximal module ahead of the impact. Note that, by necessity, the HSPC architecture includes an internal comparator that computes the prediction errors associated with the collision signal.

In keeping with the above notational conventions, the equations determining the distal feed-forward module inputs and error signals in HSPC are:

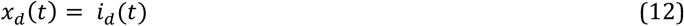

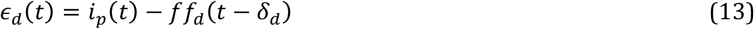

Note that the error signal that controls learning in the distal feed-forward module is a prediction error, coding the difference between a past prediction, *ff_d_*(*t* − *δ_d_*), and the actual stimulus, *i_p_(t*), where *δ_d_* is the anticipatory delay of the distal module. The proximal feed-forward module is integrated within the control architecture as follows:

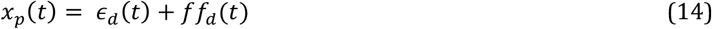

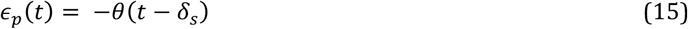

In brief, the sensory prediction error, *ϵ_d_*(*t*), and the prediction signal, *ff_d_*(*t*), related to the collision drive the proximal module, which is supervised by the error in angle (measured with a delay of *δ_s_* seconds).

In the last stage, the output of the proximal feed-forward module is added to the error in velocity driving the feedback controller. We formulate that operation by introducing *ϵ_θ_*(*t*) = −*θ*(*t* − *δ_s_*) + *ff_p_*(*t*) and then rewriting the first equation of the feedback controller:

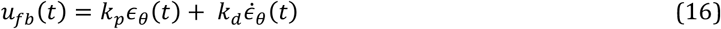

Finally, the motor control signal generated by the HSPC architecture is simply the output of the feedback controller, *u_hspc_*(*t*) = *u_fb_*(*t*).

## 3 RESULTS

Below we report on the performance of both control schemes during three experimental conditions: standard *acquisition* trials, *robustness* trials in which the disturbance is cued but not delivered, and *generalization* trials in which we provided both cued and non-cued trials, and changed the weight of the agent during training.

### 3.1.1 Acquisition

We start by analyzing the performance of the two adaptive control architectures in the acquisition of an anticipatory postural adjustment (APA) trained in a trial-by-trial manner. We use a simulated self-balancing system that at each trial receives an impact, preceded by a distal cue by a fixed interval of 400 ms, and resulting in a disturbance force (100 N during 300 ms). The force, applied to the pendulum, produces an angular displacement that, in the naive system, is uniquely counteracted by the reactive controller introducing oscillations in the angular position (Fig. 3-A, gray line). After learning, acquired motor responses evoked by the two predictive stimuli (cue and collision) substantially reduce the angular error (Fig. 3-A). Note that despite implementing different adaptive strategies, we could configure both architectures to exhibit similar learning curves (Fig. 3-B).

**Fig. 3-.**
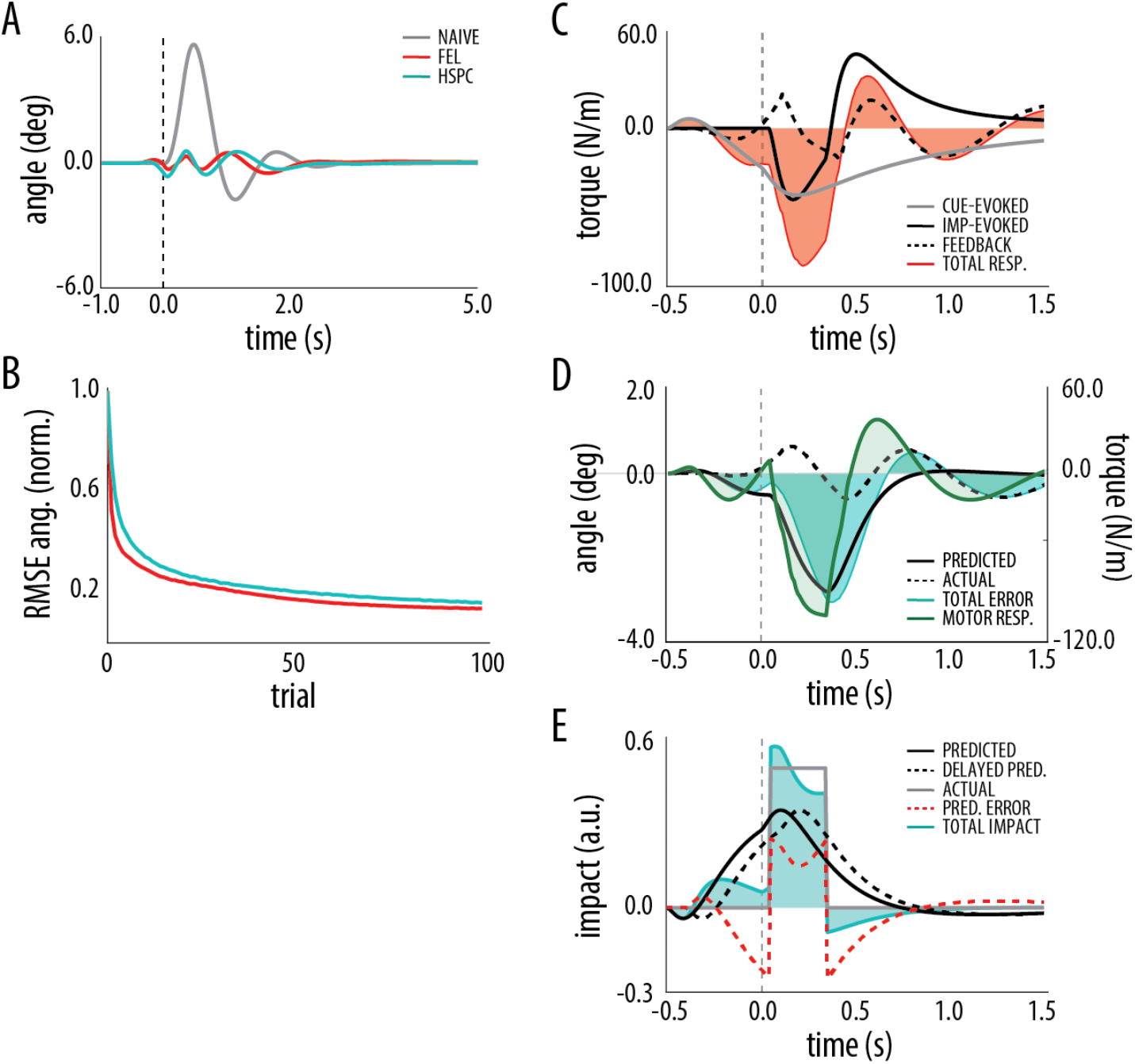
Acquisition of adaptive postural adjustment. **A.** Mean angular position during the disturbance rejection task for feedback-control condition (gray - 10 trials), trained FEL architecture (red - trials 90-100) and trained HSPC architecture (cyan - trials 90-100). Disturbance is delivered at t=0 (dashed line). **B.** Root mean square error (RMSE) in angular position over trials for FEL (red) and HSPC (cyan) architectures normalized by the maximum error in the naive system (feedback-control only). **C.** Decomposition of the motor response driving behavior in FEL: cue-evoked (gray), impact-evoked (solid black) and reactive (dashed black) responses are integrated in a total motor command (red shaded area). **D.** Decomposition of the angular error driving the behavior of the feedback controller: the total error (shaded cyan area), obtained by summing the *counterfactual* (solid black) and the current error (dashed black), is converted into a motor response (green shaded area) by the feedback controller. Error and motor response refer to different scales. **E.** Decomposition of the impact sensory signal entering the feed-forward compensatory layer of HSPC: the total impact signal (shaded cyan area) is obtained as the sum of the predicted impact signal (black solid) and the prediction error (red dashed), which is computed by subtracting the delayed prediction (black dashed) from the actual impact signal (gray solid).

After learning, in FEL the reactive controller is only marginally engaged as the errors in behavior that drove it initially are almost cancelled (Fig. 3-C). Note that in this architecture, only the cue-evoked command contributes to preparatory behavior (before the collision) but both cue- and collision-evoked commands contribute to the fast feed-forward compensation that takes place after the collision. Conversely, in HSPC the proximal adaptive module that associates the collision signal with (counterfactual) inertial errors steers the feedback controller both during anticipation and fast compensation (Fig. 3-D). Still, after learning, the proximal module is fed with a mixture of actual and anticipated collision signals, where the former is sensed and the latter provided by the distal module (Fig. 3-E). Importantly, the distal module predicts the collision signal from the cue and issues an anticipated impact signal preceding the actual impact by 100 ms (the extent of the anticipation, *δ_d_,* is a design parameter (see Methods)). Hence, the anticipatory part of the response, despite being evoked only by the cue stimulus, results from a cascade of predictions that involves both adaptive feed-forward modules and the feedback controller.

In sum, despite the marked differences in the processing, both architectures converged to similar motor commands and behavior, indicating that both motor anticipation- (FEL) and sensory prediction-based (HSPC) strategies can be equally successful in acquiring APAs.

### 3.1.2 Robustness

Next we assess the reaction of both architectures to violations in the sequence of predicted events that was learned during training. To that end, after 100 acquisition trials, we run 50 trials within which we randomly intersperse a 10% catch trials in which we present the cue but omit the disturbance. During catch trials the agent initiates an anticipatory motor response that later, due to the lack of disturbance, results in a performance error (8,20). Here, we use such error to quantify how reactive FEL and HSPC are to unexpected contingencies.

Prior to the expected impact time, both architectures introduce a slight anticipatory angular error (Fig. 4, A) by issuing the preparatory part of the response (Fig. 4B). However, once the impact fails to occur, HSPC promptly cancels the initial error while in FEL the error keeps increasing. In terms of performance, the error in a catch trial incurred by HSPC (median of the RMSE) is approximately half of the error introduced by FEL (0.3 vs 0.6 in normalized RMSE) (Fig. 4, C).

**Fig 4-.**
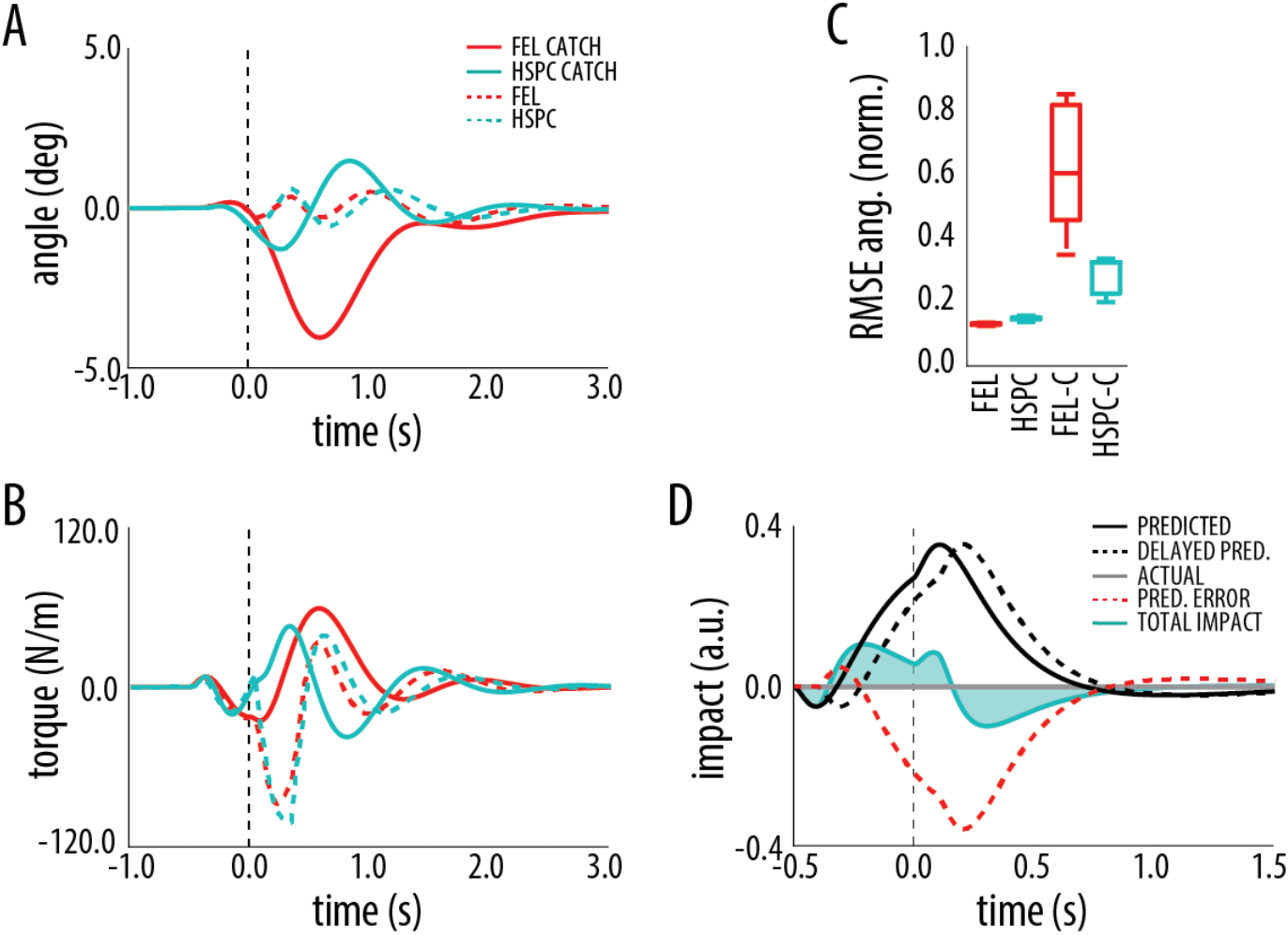
Robustness of the FEL and HSPC architectures. **A.** Mean angular position of FEL (red) and HSPC (cyan) during catch (N=10 - solid) and regular perturbed trials (N=10 - dashed). **B.** Mean motor response during catch and regular perturbed trials. Color-code as in A. **C.** Root mean square error (RMSE) in angular position during regular trained perturbed trials (N=40) and catch trials (N=10). **D.** Online prediction error correction in HSPC: the prediction error (red dashed), obtained as the difference between the delayed prediction (black dashed) and the actual impact signal (gray solid) is subtracted from the erroneously anticipated impact signal (black solid) and generates a total response (cyan area).

The reasons behind the difference in performance are the following: FEL reacts to the absence of the impact by omitting the collision-evoked command, but it issues the whole cue-evoked command (which outlasts the time of the expected collision). In contrast, HSPC rapidly aborts the compensatory action once the proximal module receives the sensory prediction error triggered by the missed collision (Fig 4, D).

In summary the HSPC architecture outperforms the FEL in that, due to the computation of sensory prediction-errors, it can react on-line to violations in the course of expected events.

### 3.1.3 Generalization

In a final set of simulations, we test how both architectures respond to changes in the plant dynamics and task contingencies. We run an additional set of 60 trials after acquisition. During the first 10 extra trials we measure the performance of the feed-forward compensatory layer in isolation, omitting the cue. At trial 11 the plant is made heavier (+10% - *light-to-heavy condition*) (note a similar manipulation in behavioral postural control studies (52)) and the agent receives additional non-cued collisions (40 trials). Afterwards, we reintroduce the cue for 10 additional trials. In a separate set of simulation, we trained initially the heavier agent and afterwards remove the weight (-10% - *heavy-to-light/condition*).

In FEL, any change in the task contingencies deteriorates performance (removing or reinstating the cue), irrespectively of whether the plant has increased or decreased its weight. In HSPC, the performance deteriorates, albeit to a lesser extent, after removing the cue. However, once the cue is reintroduced after having retrained the compensatory module, we observe a gain in performance in both cases, greater when transitioning to the lighter plant (Fig. 5-E,F).

**Fig. 5-.**
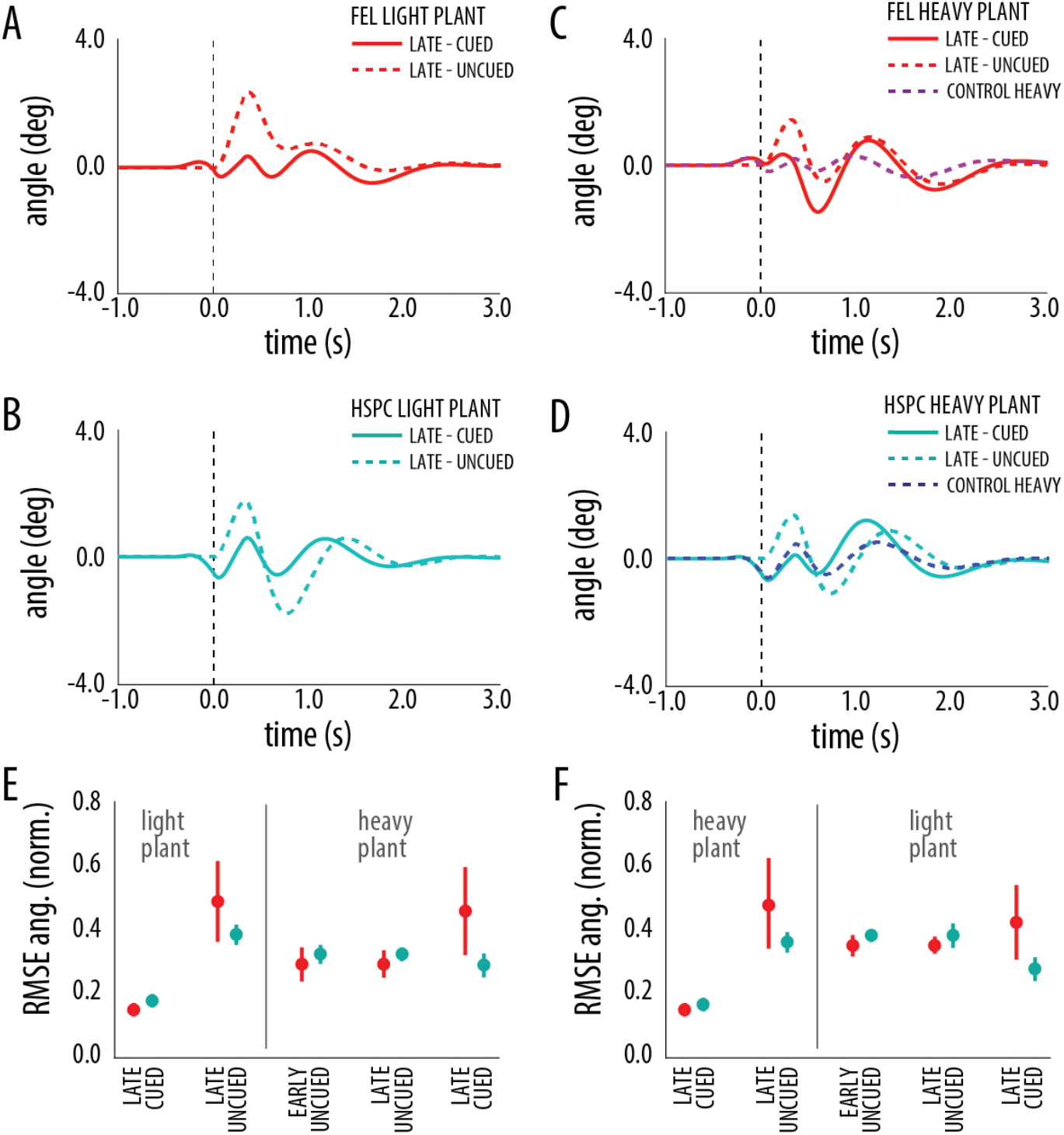
Generalization. A. FEL: Mean angular position before plant perturbation (N=10 - light plant) with (solid) and without the cue (dashed) **B. HSPC:** Same description as A. **C. FEL:** Mean angular position after plant perturbation (heavy plant - N=10) without (dashed) and with the cue (solid) and after regular training with heavy plant (solid magenta). **D. HSPC** Same description as in C. **E.** Root mean square error (RMSE) in angular position during *light-to-heavy* generalization phases for FEL (red) and HSPC (cyan). “Light plant” denotes the phase before plant perturbation. “Heavy plant” denotes the phase after plant perturbation. **F.** Same description as E for *heavy-to-light* generalization.

The difference in performance stems from the different ways in which both architectures combine the two stimuli. FEL deals with the cue and impact as independent stimuli. Initially, both contribute to the response, but once the cue is removed a part of the response is removed as well, damaging performance (Fig. 5-A). Further training makes FEL able to trigger appropriate compensatory responses just with the proximal stimulus, but then, reinstating the cue superposes a motor command partly redundant, damaging performance again (Fig. 5-C). Notably, considering that cue and impact make a compound stimulus in regular trials, the Rescorla-Wagner model (53) would describe the interference between the cue and the impact stimuli observed in FEL. On the contrary, in HSPC the distal module learns to predict the impact from the cue, and uses that prediction to trigger (a part of) the compensatory action in anticipation (Fig. 5-B). That implies that even after changing the properties of the plant, anticipating an appropriate compensatory action can result in an improvement in performance (Fig. 5-D).

In summary, in face of perturbations to the plant dynamics or changes in the task contingencies, a control strategy learning a cascade of sensory predictions allows for better generalization than one that treats the different stimuli independently.

## 4 DISCUSSION

Even though it is clearly established that skilled motor behavior relies on internal models, their nature is still under debate. The two prevailing views are that internal models can either be inverse models, mapping the desired sensory consequences into their required motor commands, or forward models, mapping motor commands into their predicted consequences. Here, we have challenged this dichotomy and advanced an alternative proposal that reformulates anticipatory motor control as a sensory-sensory learning problem (Hierarchical Sensory Prediction Control). On this view, the predicted (proprioceptive) consequences of responses to (exteroceptive) cues prescribe action or motor commands (that are mediated – or realised – by reflexes). This simplification and generalisation of the ‘standard model’ appeals to active inference; with an emphasis on estimating and predicting states of the world and the self. In order to test this hypothesis, we designed two control architectures that adopted either a motor anticipation- or a sensory prediction-based approach. We based the motor-anticipation architecture on the well-established Feedback Error Learning (FEL) model (15,16,26) whereas Hierarchical Sensory Predictive Control (HSPC) provided the sensory prediction-based architecture.

We compared both architectures in a simulated anticipatory postural adjustment (APA) task (6,8,43). Despite differences in the processing, both architectures acquired an APA equally well (Fig. 4). However, as soon as we extended the basic APA protocol with either the introduction of catch trials or by perturbing the plant, the sensory-prediction strategy outperformed motor-anticipation. Below, we will argue that the reasons for that superior performance are grounded in two specific consequences of the sensory-prediction strategy: its reliance on sensory prediction errors, and second, that HSPC affords a hierarchical processing architecture that encapsulates learning at different levels. In other words, in line with active inference, placing a hierarchical model on top of reflexive sensorimotor control equips behaviour with a context-sensitivity and intentional aspects that are precluded in ‘standard’ formulations.

### 4.1 Origin of the robustness and generalization capabilities in HSPC

The hierarchical structure of the HSPC explains its superior generalization ability. The FEL architecture has a flat structure as long as controlling behavior is concerned: all modules send motor commands in parallel to the plant. This means that after perturbing the plant, the output of all modules has to be retrained to the new plant dynamics. HSPC solves the control problem partitioning it into two smaller sub-problems: predicting the collision from the cue and predicting the postural errors from the collision. Changing the mass of the agent only changes the sensory consequences of the collision, hence, once a new feed-forward reaction to the collision is acquired, a gain in performance can still be obtained by rightly anticipating the collision (thereby, bringing the trained reaction forward in time).

On the other hand, sensory prediction errors (SPEs) enable fast reaction to erroneous predictions. As FEL only learns to react to stimuli, but not to predict them, it cannot (at least naturally) incorporate SPEs. On the contrary, HSPC relies on SPEs both for improving prediction accuracy and to preclude reaction to predicted stimuli at the time of their actual occurrence (13). That is, SPEs are intrinsic to the design principle behind HSPC. In catch trials, as no collision occurs, the prediction of the distal module fails, generating a negative SPE that interrupts the ongoing response of the proximal module initiated by the distal module, thereby enabling a fast recovery (in addition to readjustment –learning– as the absence of the collision may imply a lasting change in task contingencies).

### 4.2 Environmental forward models and inverted sensory-sensory forward models

The distal module in HSPC is a forward model of the environment that solves the problem of predicting one stimulus (a collision) given another stimulus (a cue), i.e. a task contingency. In general, forward models of the environment have been acknowledged (54), but usually not considered specifically in the context of physiological motor control except, recently, within the domain of active inference (37,40,41,55). However, the forward model in HSPC is not generically predicting one stimulus from another; it is anticipating a stimulus with the objective of driving a behavioral response that minimizes a defined error. For that, it must take into account not only sensorimotor latencies, but also the dynamics of the plant (e.g., musculo-skeletal system). Hence, the environmental forward model in HSPC affords *action-aware* predictions in that it relates external events in the context of acting to reduce a downstream error in performance.

On the other hand, the internal model dealing with the collision signal acts as an inverse model that, instead of learning postural errors, learns *from* postural errors. Its goal is not learning what the postural error signal was in a particular trial (e.g., the first one), but converging to a *counterfactual* error signal that will minimize the errors in performance. We termed this strategy counterfactual predictive control (CFPC) (33). The goal of CFPC is acquiring *counterfactual* error signals that, even though they do not code any forthcoming errors derived from the interaction with the real world, they steer a feedback controller in order to avoid actual errors to occur. In practice, this leads the adaptive model within the HSPC architecture to acquire an *inverse* model of the closed-loop system that reflects jointly the dynamics of plant and controller (33). This demonstrates how a learning process that depends on sensory errors (in contrast to motor errors) is not automatically building a forward model (e.g., (56)).

### 4.3 Related research in experimental psychology and predictions of the HSPC hypothesis

Experimental APA protocols include standing human participants receiving the impact of an object attached to a pendulum (43,45,57). As expected, those experiments show that faced with the incoming pendulum, participants rely on distal sensing (vision) to issue the anticipatory responses (43,45), i.e.: no anticipatory responses were observed when participants closed their eyes. Regarding the interplay between proprioceptive and vestibular information, separate studies in compensatory postural control have shown that humans with compromised proprioception display compensatory responses delayed with respect to healthy controls (58) as well as animals with Pyridoxine-induced loss of peripheral sensory efferents have delayed compensatory responses and increased postural sway (59). This suggests that the design of the task and the interplay between sensory modalities and responses in our simulated APA task is close agreement with biology. However, those findings do not discriminate between the sensory prediction and motor anticipation hypotheses. An exception comes from experiments showing that altered proprioceptive information at the level of the Achilles tendon delays *anticipatory* postural responses (45). Note that FEL would predict that decreasing the information in the proprioceptive channel would have no effect in the preparatory actions, which are motor commands triggered by the visual stimulus. However, in the HSPC hypothesis, anticipatory actions are elicited by generating proprioceptive predictions. Hence, one could expect that a manipulation that alters the processing of *real* proprioceptive information would also affect the mapping of predicted proprioception into action.

HSPC further predicts that in catch trials, subjects will correct erroneous anticipatory actions with latency equal to the time needed to raise sensory prediction errors. In contrast, as FEL makes no use of sensory prediction errors, there is no mechanism that could sustain such a sharp change in behavior at the expected time of the disturbance. We find evidence for such type of rapid reversals in smooth pursuit studies (27), wherein during catch trials the correction onsets at the same moment as the reactive response was triggered in early trials (i.e. sensory feedback latency) (27).

Finally, generalization of adaptive motor responses has been found in limb (28,60) and postural control (29,30). Subjects trained to catch a ball with one arm, perform equally as good when they switch arm (60), a result that cannot be explained in terms of inverse models (by definition, effector-specific). Additionally, subjects that learned to counter a force-field perturbation in a sitting position correctly anticipated the postural disturbances that compensating for the force field would introduce in an upright posture (29). This result argues in favor of an architecture composed by a forward internal representation of the dynamics of the environment coupled with an internal model of the postural dynamics, where the former is effector independent and the latter is already fine-tuned by experience; a proposal consistent with the hierarchical structure of HSPC.

Put together, these three sources of evidence (generalization of acquired responses across limbs and postures, rapid reversal of the erroneous response in catch trials, and anticipatory responses affected by altered proprioception) support a hierarchical control architecture that acquires forward models of the environment, exploits of sensory prediction errors, and shows a dependency between anticipatory and compensatory responses. All these features are embodied in HSPC but are difficult to reconcile with an inverse model-based architecture such as FEL.

### 4.5 Implications for cerebellar physiology

HSPC advances a hypothesis of cerebellar function in the domain of anticipatory control. It has its origins in a model of the cerebellum (33,48) as is the case for FEL (16). In both architectures, adaptive modules are implemented as adaptive-filters, a widely used computational model of cerebellar function (61,62). Moreover, here we have demonstrated HSPC in a task that depends on the cerebellum (63). The distinctive trait of our implementation of the cerebellar algorithm is the use of a delayed eligibility trace (Methods - Eq. 1) (48). Taking into account that in the cerebellum contextual information reaches Purkinje cells through the parallel fibers whereas specific error signals arrive via the climbing fibers, in terms of cerebellar physiology, the eligibility trace mechanism predicts a plasticity rule in the synapses between parallel fibers to Purkinje cells synapses modifies synaptic weights whenever activity in the parallel fibers precedes climbing fiber input by a certain time interval. Both in HSPC and FEL, we set that interval according the behavioral constraints of the agent/task (33), a requirement that seems to apply also in the cerebellum, where the timing of the plasticity rule of cerebellar Purkinje cells is matched to behavioral function (64). In sum, there is agreement between the requirements of the HSPC architecture and known properties of the cerebellum. In consequence, we predict that in the acquisition of anticipatory reflexes, the output of the cerebellum should be seen as sensory signals shifted forward in time, and not as anticipated motor commands as such, i.e., defining predicted error signals for the downstream targets of the cerebellum such as the Red Nucleus.

### 4.6 Summary

We have shown how a hierarchical control architecture based in the acquisition of sensory predictions enables the acquisition of reactive and generalizable APAs better than one based on the acquisition of sensory-motor associations. In doing so, we dissolved the standard inverse-forward model dichotomy by showing how (the inversion of) forward models that acquire sensory-sensory associations can contribute to motor behavior with *action-aware* sensory predictions. Our results read as a validation of key principles behind the active inference theory of motor behavior. We expect the HSPC architecture to allow for the advancement of our understanding of the mechanisms underlying physiological motor control, which we propose can now be treated in a unified active inference based framework, while also contributing to the development of robust control architectures for artificial systems.

